# The Protective Role of Intestinal Alkaline Phosphatase in Inflammatory Bowel Disease-Associated Non-Alcoholic Fatty Liver Disease

**DOI:** 10.1101/2025.02.11.637679

**Authors:** Yang Liu, Hao Lv, Haonan Zhang, Shou Yu, Qiao He, Gang Cao

## Abstract

**Background:** Inflammatory bowel disease (IBD) is a chronic and progressive inflammatory condition characterized by weight loss as a prominent feature. Non-alcoholic fatty liver disease (NAFLD), typically linked to obesity and metabolic dysregulation, is increasingly recognized as being influenced by the gut-liver axis. Notably, IBD patients exhibit a heightened susceptibility to NAFLD, although the underlying mechanisms remain poorly understood. Intestinal alkaline phosphatase (IAP), an endogenous enzyme, plays a critical role in preventing intestinal bacterial translocation. We hypothesized that IAP may serve as a potential therapeutic agent for mitigating IBD-associated NAFLD.

**Methods:** An IBD model was established using three cycles of 2% dextran sulfate sodium (DSS) administration. Mice were subsequently treated with L-phenylalanine or IAP. The activity of stool IAP, gut microbiota composition, hepatic lipid accumulation, inflammatory markers, and gut microbiome diversity were assessed.

**Results:** DSS treatment markedly reduced IAP levels. Suppression of IAP significantly increased gut permeability and exacerbated hepatic inflammation and lipid deposition. Conversely, IAP supplementation restored these parameters, improved gut microbial diversity, and normalized microbiota composition. However, IAP failed to ameliorate hepatic inflammation and lipid accumulation in Toll-like receptor 4 (TLR4) knockout mice.

**Conclusion:** Deficiency in endogenous IAP contributes to the onset of NAFLD in the context of IBD. Oral IAP supplementation enhances gut barrier integrity, stabilizes gut microbiota, and prevents NAFLD development in IBD through a TLR4-dependent mechanism.

## 1. Background

Inflammatory bowel disease (IBD) is a chronic progressive inflammatory disorder of unknown etiology, encompassing ulcerative colitis (UC) and Crohn’s disease (CD). Recently, its global incidence has been on the rise^1^. Patients with UC often present with extraintestinal manifestations, including lesions affecting the skin, joints, eyes, liver, and lungs, among which hepatic fatty lesions are the most common liver disease complication^2^.

Non-alcoholic fatty liver disease (NAFLD) is understood to result from a combination of genetic, dietary, and immunomodulatory factors and gut derived microbiotaa is considered to contribute^3-5^. Traditionally, NAFLD is thought to be more prevalent in populations with obesity and metabolic disorders^3^. However, IBD is generally considered a wasting disease, characterized in some cases by malabsorption and significant weight loss^6^. Recent studies utilizing various imaging modalities have shown a higher prevalence of NAFLD in UC patients compared to the general population^7^. The mechanisms underlying the occurrence of NAFLD in IBD patients remain unclear. Research has identified that the development of NAFLD is closely associated with lifestyle factors, and its pathogenesis is multifactorial, involving genetic susceptibility, gut microbiota composition, and insulin .resistance, among other interactions^8, 9^.

Intestinal alkaline phosphatase (IAP) is one of four isoenzymes of alkaline phosphatase discovered in recent years, the others being tissue nonspecific alkaline phosphatase (found in liver, bone, and kidney), placental alkaline phosphatase, and germ cell alkaline phosphatase^10^. IAP is uniquely expressed in the intestine and produced by intestinal epithelial cells; it is a glycoprotein anchored to these cells, released into the intestinal lumen postprandially to exert its effects^10^. IAP has been found to detoxify LPS and stabilize gut microbiota. Our previous studies indicated that reduced IAP levels contribute to metabolic syndrome and age-related intrahepatic fat deposition^11-13^. Additionally, we found that IAP can lower LPS concentrations in the portal venous system, improving gut-derived hepatic inflammation^14, 15^. Based on this, we speculate that IAP may have a significant correlation with IBD-related NAFLD.

## 2. Methods

### 2.1 Animal Model

Male C57BL/6L mice aged 6-7 weeks were acclimatized for one week before being grouped for modeling. Mice received three cycles of 2% DSS, with continuous access to 2% DSS for 7 days in each cycle, replacing DSS every two days, followed by a 7-day period with drinking water. Animals were euthanized 24 hours after the third cycle, and samples were collected for analysis.

### 2.2 Oral IAP or L-phenylalanine

IAP (500 IU) or IAP specific inhibitor L-phenylalanine (16mM) was daily gavaged, while equal volume of PBS was gavaged in control group. IAP administration began 4 days prior to DSS treatment and continued throughout the experiment, with daily replacement of IAP-supplemented drinking water.

### 2.3 Statistical Analysis

Data analysis was performed using GraphPad Prism 8.0 software. Numerical data are expressed as mean **±** standard deviation. Differences between groups were compared using Student’s t-test. Pearson correlation analysis was conducted for serum LPS and MPO-DNA. A p-value of less than 0.05 was considerd statistically significant.

Detailed methods are described in the supplementary methods.

## 3. Results

### DSS treatment reduces intestinal IAP activity, impairs intestinal barrier integrity, and promotes hepatic fat accumulation

We initially assessed stool IAP activity using pNPP as a substrate to measure IAP activity. After three cycles of DSS treatment, we observed a significant reduction in IAP activity in the feces compared to sham treatment, accompanied by a decrease in the expression of IAP-related genes, AKP3 and AKP6, in the duodenum of mice (Fig. 1A). Concurrently, the expression of tight junction related genes ZO1, ZO2, and Occludin in colon, as assessed by qPCR, as well as the positive staining area revealed by immunohistochemical analysis, was significantly reduced following DSS treatment (Fig. 1C, D). Additionally, serum levels of LPS and FITC-dextran, both indicators of compromised gut barrier function, were significantly elevated in the DSS-treated group compared to sham group(Fig. 1E, F).

**Figure 1.**
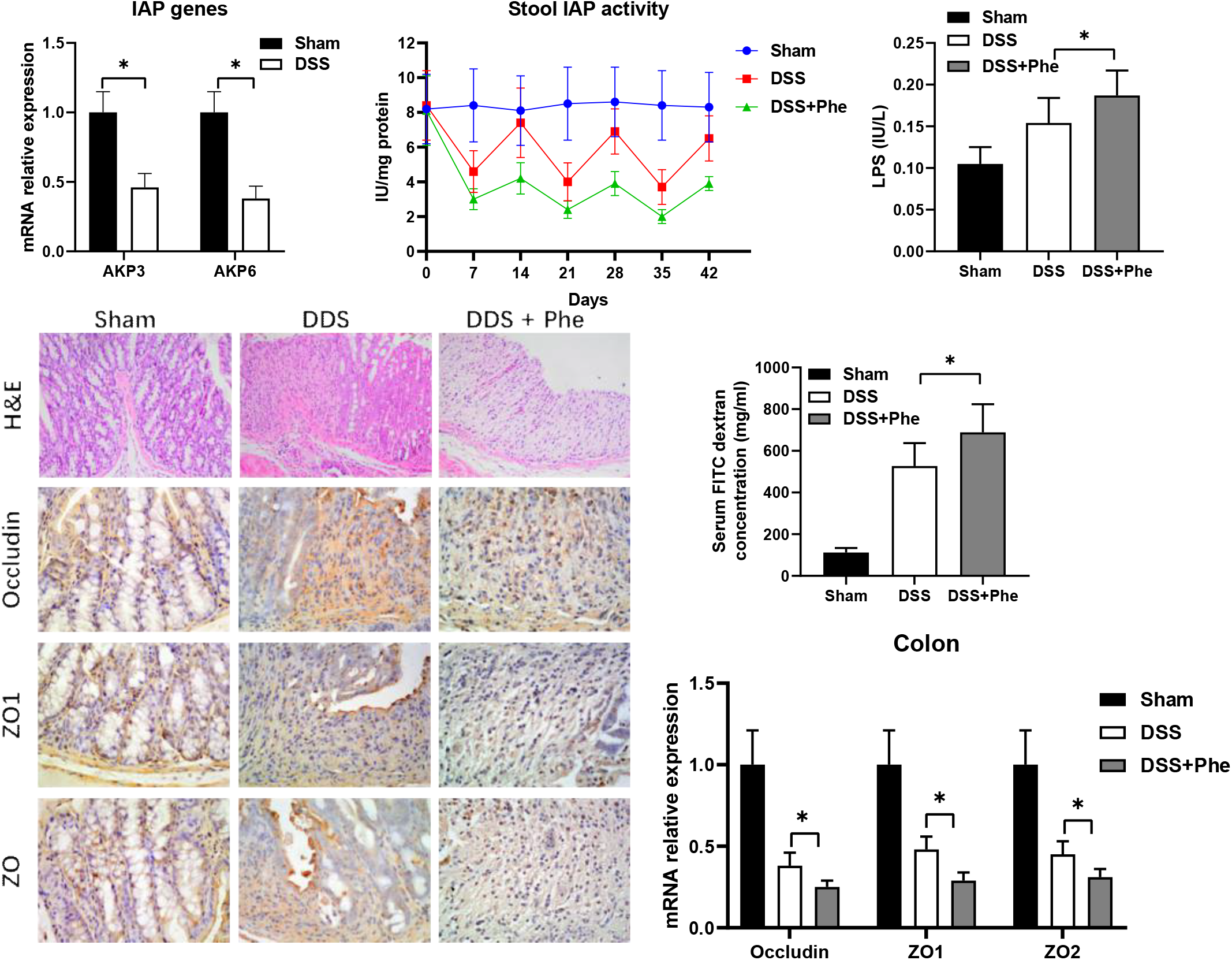
DSS treatment reduces intestinal IAP activity, and IAP deficiency results in severe intestinal barrier damage. (A) Expression of IAP-related genes in DSS-treated versus sham-treated mice, as measured by qPCR. (B) Stool IAP activity measured using the pNPP assay. (C) Immunohistochemical staining of Occludin, ZO1, and ZO2 in the colon following DSS or DSS + L-phenylalanine treatment. (D) Serum LPS concentrations in DSS- and DSS + L-phenylalanine-treated mice, measured using a detection kit. (E) Serum FITC-dextran levels 4 hours post-gavage in the indicated groups.(F) Gene expression of Occludin, ZO1, and ZO2 in the mouse colon, determined by qPCR for each group. *p < 0.05.

Subsequently, we evaluated hepatic triglyceride (TG) concentrations, which were found to be significantly higher in the DSS-treated group than sham group(Fig. 2B). Similar results were confirmed by Oil Red O staining of liver tissues (Fig. 2D). Furthermore, the expression of inflammatory markers in the liver, including TNFα, IL6, and IL1β, was markedly increased, while genes associated with fat oxidation and degradation, such as ATGL, MGL, HSL, PGC1α, CPT1, and MCAD, were significantly downregulated following three cycles of DSS treatment than sham treatment(Fig. 2C, E).

**Figure 2.**
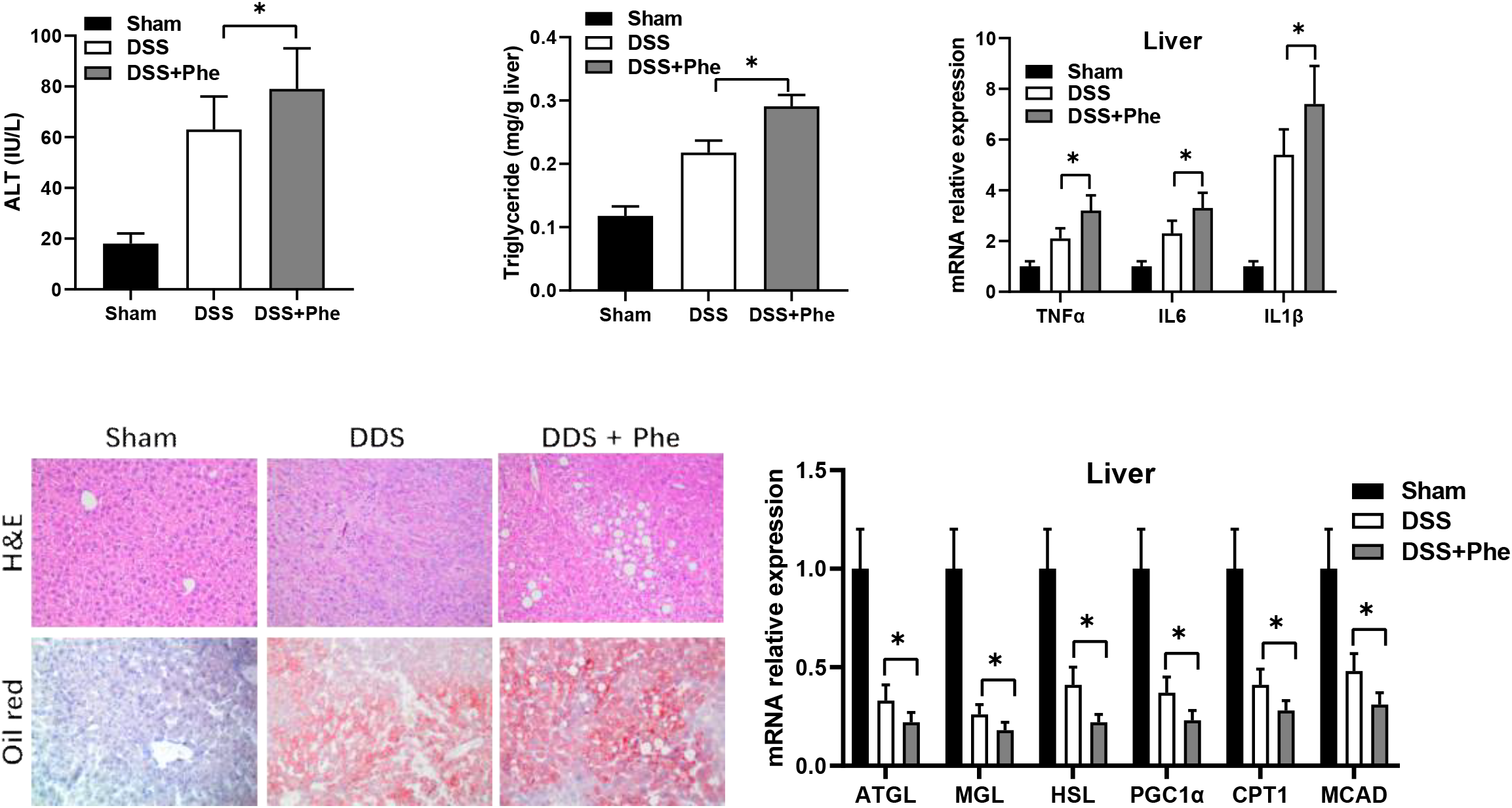
Inhibition of IAP exacerbates inflammatory responses and promotes fat deposition in the liver. (A) Serum ALT levels in DSS- and DSS + L-phenylalanine-treated mice, measured by a detection kit. (B) Liver triglyceride concentrations in different treatment groups, revealed using a detection kit. (C) Expression of liver inflammation-related genes (TNFα, IL6, and IL1β) in DSS- and DSS + L-phenylalanine-treated mice, determined by qPCR. (D) Representative liver sections stained with H&E and Oil Red O. (E) Expression of fat deposition-related genes (ATGL, MGL, HSL, and PGC1α) in the liver, determined by qPCR. *p < 0.05.

### Lack of IAP results in a severe intestinal barrier damage

Our previous findings indicated that IAP levels decrease during DSS-induced hepatic fat deposition, and it has been established that IAP plays a crucial role in various liver lesions^13, 14^. Thus, we hypothesized that IAP may also be involved in IBD-related NAFLD. In this study, we employed L-phenylalanine, a specific inhibitor of IAP. Administration of L-phenylalanine + DSS to mice resulted in a further reduction of fecal IAP levels than DSS treatment (Fig. 1B). qPCR analysis of the ZO1, ZO2, and Occludin genes in the terminal ileum revealed significant reductions in these barrier-related markers in L-phenylalanine + DSS group than DSS group (Fig. 1C). Immunohistochemical staining of ZO1, ZO2, and Occludin in the colonic tissue showed a significant decrease in the expression of these proteins in L-phenylalanine + DSS group than DSS group (Fig. 1D). After measuring LPS and FITC-dextran in portal vein serum, we found that L-phenylalanine + DSS treatment significantly elevated serum LPS and FITC-dextran levels than DSS treatment (Fig. 1E, F).

### Inhibition of IAP leads to more severe inflammatory responses and fat deposition in the liver

Additionally, we detected a significant increase in ALT concentrations in the serum following L-phenylalanine administration (Fig 2A). qPCR analysis revealed significant increases in liver inflammatory markers TNFα, IL6, and IL1β upon L-phenylalanine + DSS treatment than DSS treatment (Fig. 2B). Histological analysis showed that L-phenylalanine + DSS treatment resulted in more extensive fat vacuole formation in the liver, and Oil Red O staining revealed more red-stained areas than DSS treatment (Fig. 2C). Using a kit to measure TG content in liver tissue yielded similar results (Fig. 2D). Further qPCR analysis of fat oxidation and degradation-related markers, such as ATGL, MGL, HSL, PGC1α, CPT1, and MCAD, indicated a decrease in these indicators associated with IAP reduction upon plus L-phenylalanine administration (Fig. 2 E).

### IAP supplementation alleviates DSS-induced intestinal barrier impairment and changes in intestinal permeability

Having established that oral IAP improves various liver disease processes by enhancing the intestinal barrier, we sought to prevent DSS-induced NAFLD through exogenous IAP supplementation. We found that mice receiving IAP + DSS treatment showed significant weight improvement compared to the DSS group (Fig. 3A). After three cycles of DSS, the serum FITC levels in IAP-treated mice were markedly lower than those in untreated mice (Fig. 3B). We further measured LPS levels in the portal vein serum of IAP-treated mice, revealing a significant reduction in LPS levels post-treatment (Fig. 3C). Immunohistochemical analysis of ZO1, ZO2, and Occludin in the colonic tissue showed significantly reduced positive staining regions after IAP treatment. Further qPCR analysis demonstrated that the expression levels of ZO1, ZO2, and Occludin genes in the terminal ileum were decreased in IAP-treated mice (Fig. 3D, E).

**Figure 3.**
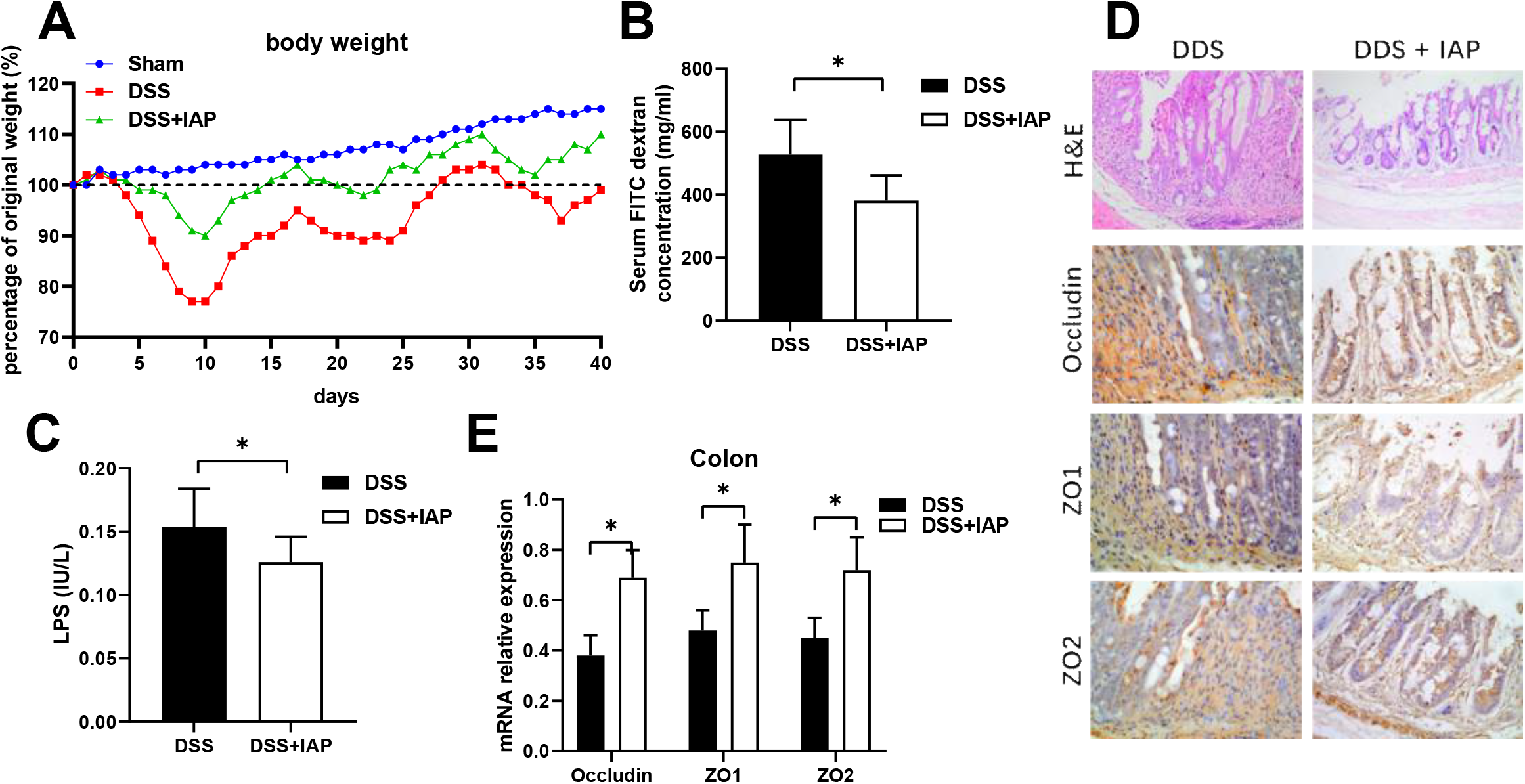
IAP supplementation mitigates DSS-induced intestinal barrier impairment and alterations in intestinal permeability. (A) Body weight changes in each group. (B) Serum FITC-dextran levels 4 hours post-gavage in the indicated groups. (C) Immunohistochemical staining of Occludin, ZO1, and ZO2 in the colon following DSS or DSS + IAP treatment. (D) Gene expression of Occludin, ZO1, and ZO2 in the mouse colon, determined by qPCR for each group. *p < 0.05.

### IAP reduces liver inflammation and lowers fatty liver

The ALT levels in serum also showed a notable decrease after plus IAP administration (Fig. 4A). The measurement of triglyceride content in liver tissue also confirmed that oral IAP significantly reduced hepatic triglyceride levels (Fig. 4B). HE staining of liver tissue showed a marked reduction in fat vacuoles following IAP + DSS treatment than DSS treatment. Oil Red O staining demonstrated a significant reduction in the area of red-stained fat in the livers of IAP-treated mice compared to controls (Fig. 4C). Further examination of the liver tissue indicated significantly reduced gene expression levels of inflammatory markers TNFα, IL6, and IL1β (Fig. 4D). qPCR analysis of genes related to fat metabolism and oxidation (ATGL, MGL, HSL, PGC1α, CPT1, and MCAD) showed that IAP improved the expression of these markers (Fig. 4E).

**Figure 4.**
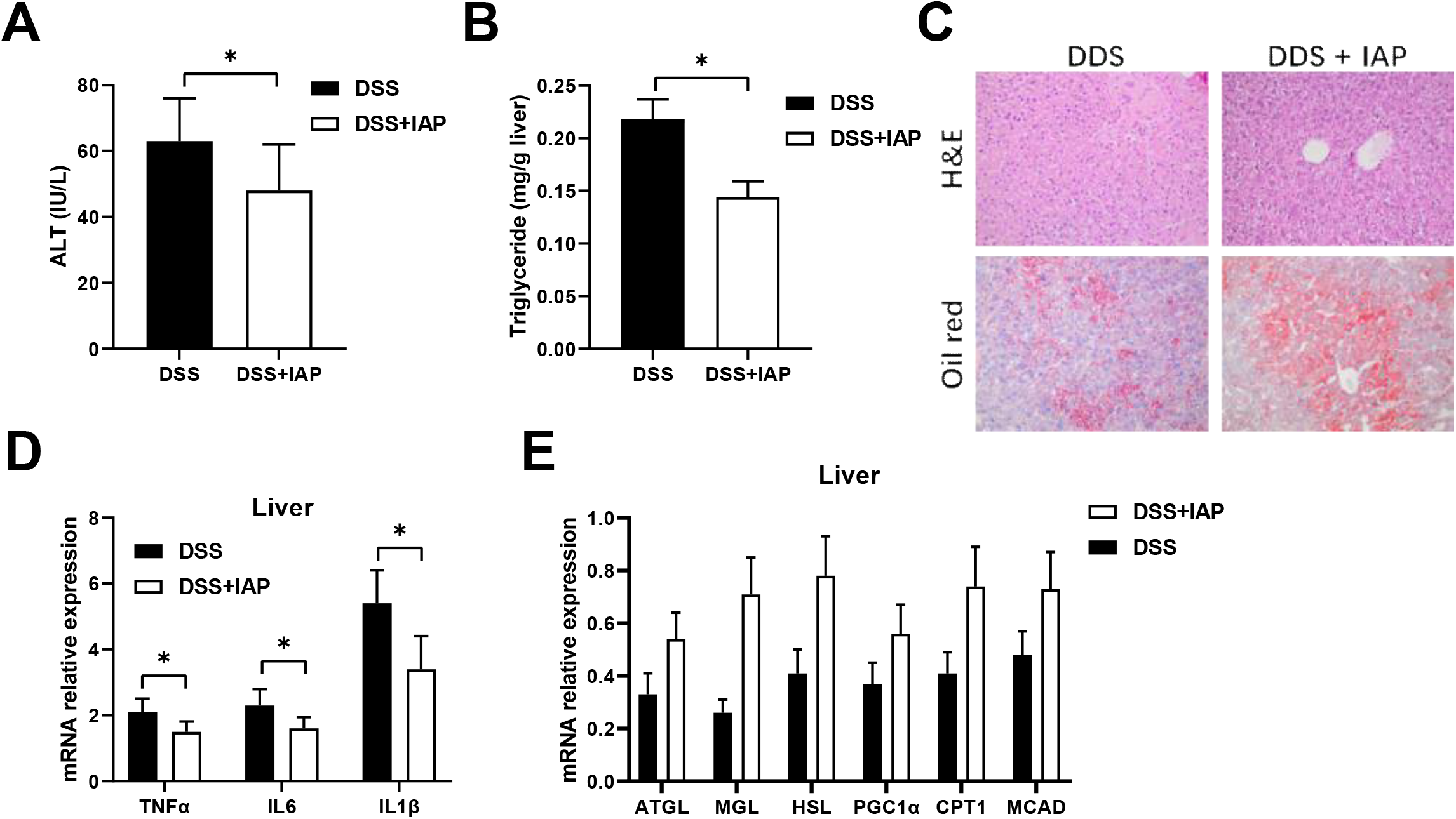
IAP reduces liver inflammation and ameliorates fatty liver. (A) Serum ALT levels in DSS and DSS + IAP-treated mice, measured by a detection kit. (B) Liver triglyceride concentrations in different groups, revealed using a detection kit. (C) Representative liver sections stained with H&E and Oil Red O. (D) Expression of liver inflammation-related genes (TNFα, IL6, and IL1β) in DSS- and DSS + IAP-treated mice, determined by qPCR. (E) Expression of fat metabolism-related genes (ATGL, MGL, HSL, and PGC1α) in the liver, determined by qPCR. *p < 0.05.

### IAP Stabilizes Gut Microbial Distribution in Mice with DSS-Induced Fatty Liver

In order to clarify the function and potential mechanism of IAP in rescue DSS induce fatty liver, we performed 16S rRNA high-throughput sequencing to analyze the gut microbiome from colonic feces. The results indicated that there were 135 shared operational taxonomic units (OTUs) among the groups. Additionally, the sham group, DSS group, and IAP group had 132, 122, and 81 unique OTUs, respectively. The sham treatment group shared 45 OTUs with the DSS treatment group and 74 OTUs with the DSS + IAP group (Fig. 5D). Regarding microbial diversity, the Shannon, Simpson, and Chao1 indices indicated a decrease in α-diversity of the gut microbiome induced by DSS, indicating significant differences in microbial composition. However, IAP treatment showed a trend towards stabilizing diversity (Fig. 5A-C). At the phylum level, the most abundant groups were Bacteroidetes, Firmicutes, Epsilonbacteraeota, Patescibacteria, and Proteobacteria, collectively accounting for over 95% of the composition. Compared to the control group, DSS-induced colitis significantly increased the relative abundances of Bacteroidetes (from 55.17% to 39.21%) and Proteobacteria (from 3.81% to 4.38%), while Firmicutes (from 16.38% to 45.84%), Epsilonbacteraeota (from 3.45% to 0.96%), and Patescibacteria (from 0.26% to 0.58%) showed significant decreases. Notably, the Firmicutes/Bacteroidetes ratio in DSS (0.87) was lower than that in the normal group (3.37), while IAP supplementation restored the Firmicutes abundance and the Firmicutes/Bacteroidetes ratio (1.41) (Fig. 5E). These results suggest that IAP effectively improves gut microbial diversity and composition in DSS-induced colitis mice.

**Figure 5.**
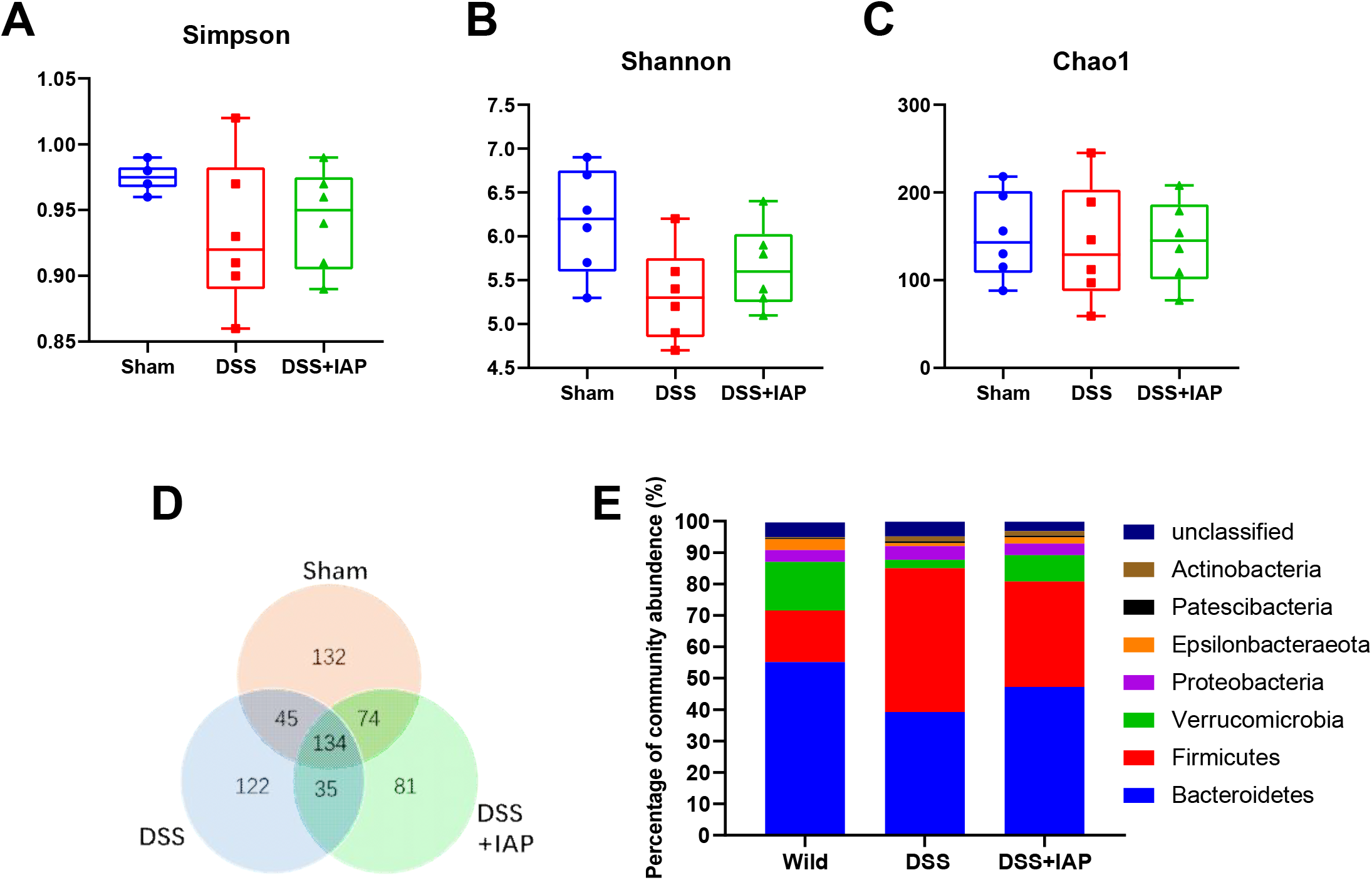
Effects of IAP on gut bacterial diversity. (A) Simpson index, (B) Shannon index, (C) Chao1 index, (D) Venn diagram showing shared and unique operational taxonomic units, and (E) Gut microbiota composition at the phylum level in the indicated groups. *p < 0.05.

### IAP rescues IBD related NAFLD dependent of TLR4 pathway

Given that IAP administration could maintaining gut microbiome homeostasis and gut barrier. Considering that gut microbiome play a crucial role in NAFLD occurrence via LPS^16^, we hypothesized that IAP may exert its effects on DSS-induced fatty liver through the LPS-TLR4 pathway. To test this hypothesis, we utilized TLR4 KO mice. In our study, serum ALT concentration and liver TG concentration was lower in TLR4 mice than WT mice after three cycle of DSS treatment (Fig. 6A, B). mRNA expressions of TNFα, IL6, IL1β, ATGL, MGL, HSL, PGC1α, CPT1, and MCAD were significantly higher in TLR4 mice compared to WT mice after DSS treatment (Fig. 6C, E). Oil red positive staining area were also significantly decreased in TLR4 KO mice compared to WT mice (Fig. 6D). This finding is consistent with previous results in murine NAFLD models.

**Figure 6.**
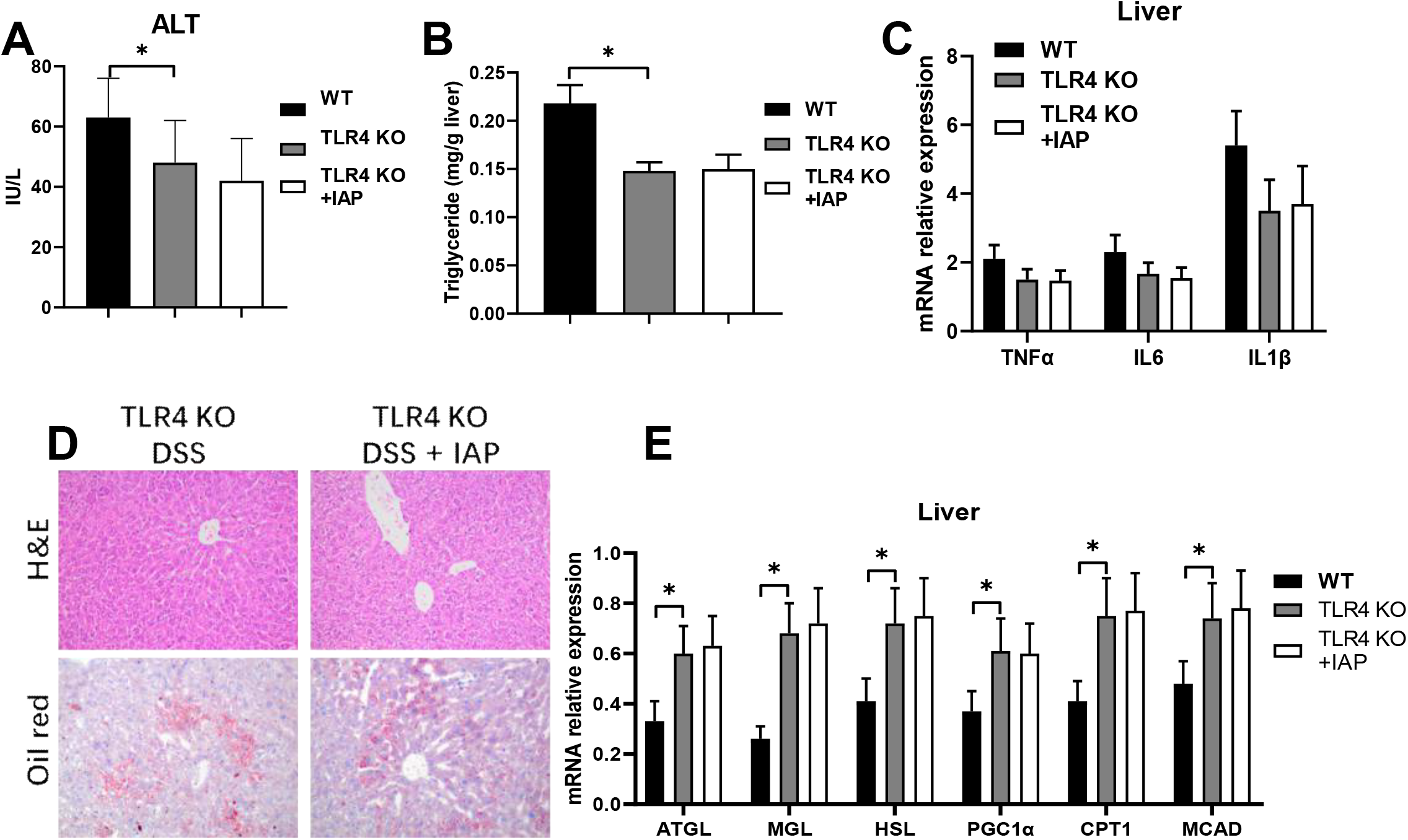
IAP ameliorates IBD-associated NAFLD via the TLR4 pathway. (A) Serum ALT levels in TLR4 KO mice treated with DSS or DSS + IAP, measured using a detection kit. (B) Liver triglyceride concentrations in TLR4 KO mice from different treatment groups, revealed using a detection kit. (C) Representative liver sections stained with H&E and Oil Red O. (D) Expression of liver inflammation-related genes (TNFα, IL6, and IL1β) in TLR4 KO mice treated with IAP, determined by qPCR. (E) Expression of fat metabolism-related genes (ATGL, MGL, HSL, and PGC1α) in the liver of TLR4 KO mice, determined by qPCR. *p < 0.05.

While it is known that LPS is a substrate for IAP, and that IAP decreases portal LPS, we asked the question if the ability of IAP to prevent IBD related NAFLD is dependent on the TLR4 pathway. Accordingly, we performed our IBD related NAFLD using TLR4 KO mice. We found that while TLR4 KO mice did not display a different serum ALT concentration and liver TNFα, IL6, IL1β, ATGL, MGL, HSL, PGC1α, CPT1, and MCAD mRNA expression than WT mice after DSS treatment between IAP supplementation and control-treated mice (Fig. 6A, C, E). Liver TG concentration was also similar between IAP or vehicle treatment in TLR4 KO mice (Fig. 5B). Similarly, Sirius oil red and HE staining also did not show a difference between IAP or vehicle-treated mice after DSS treatment (Fig. 6D).

## Discussion

According to a recent meta-analysis, NAFLD affects approximately 25% of the global population^17^. It is well-established that NAFLD is closely associated with metabolic disorders such as diabetes, dyslipidemia, and obesity. However, patients with IBD often experience malnutrition and weight loss. Studies show that the prevalence of NAFLD among IBD patients is 32%, which is higher than the prevalence in the general population as reported in meta-analyses^18^. It is generally believed that impaired intestinal mucosal barrier function, active and chronic IBD, particularly CD, small bowel resection, sarcopenia, and certain drug treatments contribute to the high incidence of NAFLD in IBD^7-9^. However, whether other factors are involved remains to be further investigated.

IAP is known as a marker of intestinal epithelial cell maturation, and its activity is negatively correlated with intestinal inflammation. IAP activity has been shown to decrease in chronic intestinal inflammatory diseases such as IBD, alcohol consumption, celiac disease, metabolic syndrome, and obesity^11, 12, 19^. The intestinal epithelial barrier is a natural barrier that absorbs nutrients and prevents harmful substances in the gut from entering the portal circulation and damaging the liver^20, 21^. Increased intestinal permeability associated with active IBD has been implicated in NAFLD, with a prevalence rate of 50-60%^22^. The integrity of the intestinal barrier is maintained by tight junction (TJ) proteins, including mucin 2, ZO-1, and occludin. Increased intestinal permeability facilitates the entry of gut-derived products such as lipopolysaccharides (LPS) and cytokines into the gut-liver circulation, exposing the liver to these byproducts^20, 21^. In this study, we found that oral IAP administration could restore the structural damage of the intestinal barrier caused by DSS. Subsequently, LPS activates toll-like receptor (TLR)-mediated inflammatory pathways, increasing the risk of NAFLD. Overall, the consequences of leaky gut may lead to liver injury, inflammation, and even NASH.

In this study, we found that IBD mice with concurrent NAFLD had lower fecal IAP levels. L-phenylalanine, as a selective IAP inhibitor, has been shown to inhibit IAP activity under various conditions^23^. Additionally, in studies involving aspartame, it was found that aspartame may inhibit IAP’s function further by generating L-phenylalanine, promoting hepatic fat deposition^24^. We observed that administration of L-phenylalanine further enhanced hepatic fat deposition in mice, providing preliminary evidence of the critical role of IAP in the promotion of NAFLD in IBD.

Moreover, fecal IAP levels are considered a potential predictor of metabolic syndromes such as diabetes^19, 25, 26^. Our study similarly found a decrease in fecal IAP levels in IBD-related NAFLD, suggesting it may serve as another simple predictor of the risk for IBD-related NAFLD. However, further research is needed to confirm whether the level of IAP decline correlates with the severity of NAFLD.

Although NAFLD is a common condition, there are currently little approved pharmacological treatments for it^27, 28^. In this study, we demonstrated for the first time that long-term oral IAP could prevent the occurrence of IBD-related NAFLD, providing a potential clinical preventive option. As an endogenous enzyme secreted by intestinal epithelial cells, IAP is theoretically non-toxic with long-term use^29^. Our previous studies have shown no adverse effects from lifelong oral IAP in mice^13^. A human study administering enteral IAP to patients with severe ulcerative colitis for seven days reported no safety issues, adverse events, or side effects^30^. Existing research indicates that IAP has been shown to improve the occurrence of extrahepatic diseases by influencing the gut-liver axis in various models, including alcoholic fatty liver, cirrhosis, and burns^12, 14, 15^. Notably, we recently demonstrated that long-term oral IAP could extend mouse lifespan, providing an inherent impetus for the potential long-term daily administration of this treatment^13^.

The gut microbiome is another key factor in gut-liver axis^21^. Various drugs, such as probiotics and butyrate formulations, have been shown to stabilize gut microbiota and inhibit the occurrence of NAFLD^31, 32^. Additionally, butyrate, as a non-selective HDAC inhibitor, has been shown to stabilize gut microbiota while promoting IAP^10^. In our study, IAP administration limited the dysbiosis induced by DSS, reducing the expression of certain bacteria while increasing the proportion of others. Gram-negative bacteria are the primary source of gut-derived toxins, and concurrent intestinal barrier damage can lead to elevated levels of LPS in the portal circulation, suggesting an increase in hepatic translocation. Furthermore, in TLR4 KO mice, IAP lost its protective effect against NAFLD. Our findings further indicate that the preventive effect of IAP against IBD-related NAFLD may likely be mediated through interventions in gut bacteria and the LPS-TLR4 pathway.

However, there are still several limitations in this study. First, while we proposed a correlation between decreased fecal IAP levels and NAFLD, the extent and duration of this decline in relation to NAFLD severity remain uncertain. Second, we began administering IAP prior to DSS treatment to intervene in the gut-liver axis to prevent NAFLD. It remains to be determined whether IAP can reverse the occurrence of NAFLD or further inhibit the progression of NASH after its onset.

In summary, our study proposes that the reduction of IAP is one of the reasons for the occurrence of NAFLD in IBD, and supplementation with exogenous IAP can effectively mitigate the incidence of DSS-induced NAFLD. This effect of IAP may be mediated through stabilizing the gut microbiota and the LPS-TLR4 pathway, providing a potential preventive strategy for IBD-related NAFLD.

## References

1. Lv H, Li HY, Zhang HN, et al. Delayed diagnosis in inflammatory bowel disease: Time to consider solutions. World J Gastroenterol 2024;30:3954–3958.

2. Rogler G, Singh A, Kavanaugh A, et al. Extraintestinal Manifestations of Inflammatory Bowel Disease: Current Concepts, Treatment, and Implications for Disease Management. Gastroenterology 2021;161:1118–1132.

3. Machado MV, Cortez-Pinto H. NAFLD, MAFLD and obesity: brothers in arms? Nat Rev Gastroenterol Hepatol 2023;20:67–68.

4. Kolodziejczyk AA, Zheng D, Shibolet O, et al. The role of the microbiome in NAFLD and NASH. EMBO Mol Med 2019;11.

5. Wang R, Tang R, Li B, et al. Gut microbiome, liver immunology, and liver diseases. Cell Mol Immunol 2021;18:4–17.

6. Barreiro-de Acosta M, Molero A, Artime E, et al. Epidemiological, Clinical, Patient-Reported and Economic Burden of Inflammatory Bowel Disease (Ulcerative colitis and Crohn’s disease) in Spain: A Systematic Review. Adv Ther 2023;40:1975–2014.

7. Ritaccio G, Stoleru G, Abutaleb A, et al. Nonalcoholic Fatty Liver Disease Is Common in IBD Patients However Progression to Hepatic Fibrosis by Noninvasive Markers Is Rare. Dig Dis Sci 2021;66:3186–3191.

8. Papaefthymiou A, Potamianos S, Goulas A, et al. Inflammatory Bowel Disease-associated Fatty Liver Disease: the Potential Effect of Biologic Agents. J Crohns Colitis 2022;16:852–862.

9. Wei Z, Wang J. Exploration of the core pathway of inflammatory bowel disease complicated with metabolic fatty liver and two-sample Mendelian randomization study of the causal relationships behind the disease. Front Immunol 2024;15:1375654.

10. Santos GM, Ismael S, Morais J, et al. Intestinal Alkaline Phosphatase: A Review of This Enzyme Role in the Intestinal Barrier Function. Microorganisms 2022;10.

11. Kaliannan K, Hamarneh SR, Economopoulos KP, et al. Intestinal alkaline phosphatase prevents metabolic syndrome in mice. Proc Natl Acad Sci U S A 2013;110:7003–8.

12. Hamarneh SR, Kim BM, Kaliannan K, et al. Intestinal Alkaline Phosphatase Attenuates Alcohol-Induced Hepatosteatosis in Mice. Dig Dis Sci 2017;62:2021–2034.

13. Kuhn F, Adiliaghdam F, Cavallaro PM, et al. Intestinal alkaline phosphatase targets the gut barrier to prevent aging. JCI Insight 2020;5.

14. Liu Y, Cavallaro PM, Kim BM, et al. A role for intestinal alkaline phosphatase in preventing liver fibrosis. Theranostics 2021;11:14–26.

15. Adiliaghdam F, Cavallaro P, Mohad V, et al. Targeting the gut to prevent sepsis from a cutaneous burn. JCI Insight 2020;5.

16. Ohtani N, Kamiya T, Kawada N. Recent updates on the role of the gut-liver axis in the pathogenesis of NAFLD/NASH, HCC, and beyond. Hepatol Commun 2023;7.

17. Lv H, Liu Y. Management of non-alcoholic fatty liver disease: Lifestyle changes. World J Gastroenterol 2024;30:2829–2833.

18. Lin A, Roth H, Anyane-Yeboa A, et al. Prevalence of Nonalcoholic Fatty Liver Disease in Patients With Inflammatory Bowel Disease: A Systematic Review and Meta-analysis. Inflamm Bowel Dis 2021;27:947–955.

19. Malo MS. A High Level of Intestinal Alkaline Phosphatase Is Protective Against Type 2 Diabetes Mellitus Irrespective of Obesity. EBioMedicine 2015;2:2016–23.

20. Luo L, Chang Y, Sheng L. Gut-liver axis in the progression of nonalcoholic fatty liver disease: From the microbial derivatives-centered perspective. Life Sci 2023;321:121614.

21. Pabst O, Hornef MW, Schaap FG, et al. Gut-liver axis: barriers and functional circuits. Nat Rev Gastroenterol Hepatol 2023;20:447–461.

22. Likhitsup A, Dundulis J, Ansari S, et al. High prevalence of non-alcoholic fatty liver disease in patients with inflammatory bowel disease receiving anti-tumor necrosis factor therapy. Ann Gastroenterol 2019;32:463–468.

23. Jassas RS, Naeem N, Sadiq A, et al. Current status of N-, O-, S-heterocycles as potential alkaline phosphatase inhibitors: a medicinal chemistry overview. RSC Adv 2023;13:16413–16452.

24. Gul SS, Hamilton AR, Munoz AR, et al. Inhibition of the gut enzyme intestinal alkaline phosphatase may explain how aspartame promotes glucose intolerance and obesity in mice. Appl Physiol Nutr Metab 2017;42:77–83.

25. Malo J, Alam MJ, Islam S, et al. Intestinal alkaline phosphatase deficiency increases the risk of diabetes. BMJ Open Diabetes Res Care 2022;10.

26. Lassenius MI, Fogarty CL, Blaut M, et al. Intestinal alkaline phosphatase at the crossroad of intestinal health and disease - a putative role in type 1 diabetes. J Intern Med 2017;281:586–600.

27. Ferguson D, Finck BN. Emerging therapeutic approaches for the treatment of NAFLD and type 2 diabetes mellitus. Nat Rev Endocrinol 2021;17:484–495.

28. Ciardullo S, Muraca E, Vergani M, et al. Advancements in pharmacological treatment of NAFLD/MASLD: a focus on metabolic and liver-targeted interventions. Gastroenterol Rep (Oxf) 2024;12:goae029.

29. Kuhn F, Duan R, Ilmer M, et al. Targeting the Intestinal Barrier to Prevent Gut-Derived Inflammation and Disease: A Role for Intestinal Alkaline Phosphatase. Visc Med 2021;37:383–393.

30. Lukas M, Drastich P, Konecny M, et al. Exogenous alkaline phosphatase for the treatment of patients with moderate to severe ulcerative colitis. Inflamm Bowel Dis 2010;16:1180–6.

31. Sarkar A, Mitra P, Lahiri A, et al. Butyrate limits inflammatory macrophage niche in NASH. Cell Death Dis 2023;14:332.

32. Arai N, Miura K, Aizawa K, et al. Probiotics suppress nonalcoholic steatohepatitis and carcinogenesis progression in hepatocyte-specific PTEN knockout mice. Sci Rep 2022;12:16206.

